# Evolution and information content of optimal gene regulatory architectures

**DOI:** 10.1101/2025.06.10.657849

**Authors:** Nicholas H. Barton, Gašper Tkačik

## Abstract

Evolution of gene regulatory sequences is shaped by their genotype-phenotype (GP) map, but this map itself can evolve via changes in the molecular machinery that reads out the regulatory sequences. Using population genetics theory, we derive an optimality principle, implying that the evolved GP maps will associate fit phenotypes with larger numbers of possible genotypes. Mathematically, this builds on analogies between population genetics and statistical physics, as well as optimal coding in information theory. The results are particularly interesting in the context of transcriptional regulation, where optimal values of certain parameters that determine the gene regulatory architecture can be derived even without a complete knowledge of the molecular mechanisms involved. We illustrate the theory using a simplified model of transcription factor (TF) binding to cis-regulatory elements (CREs): the fraction of possible CREs across the entire genotype space that bind a given TF evolves to match the fraction of CREs under selection to bind that TF. Similar “frequency matching” results are expected to emerge generically, because selection implicitly optimizes the entire regulatory architecture that determines the “replicated GP map” for all CREs, while it simultaneously adapts towards individually fit CREs on that GP map. We discuss the subtleties and limitations of the theory related to biophysical constraints and the need for mutational robustness. Lastly, our theory suggests a possible mathematical definition for evolvability with predictive power that can be tested in evolutionary simulations and, ultimately, in comparative genetics studies.

---

Evolving genomes experience selection that is shaped by the underlying GP map. For example, regulatory sequences such as promoters or enhancers may be selected to bind specific TFs. But the GP map itself can evolve: changes to the TF concentration or the TF’s DNA-binding domain will affect which cis-regulatory element (CRE) sequences bind that TF, via changes to the size, sequence, and number of potential binding sites. Mutations that affect the map may have highly pleiotropic effects, because a single TF can interact with CREs scattered across the entire genome, and therefore experience strong selection associated with phenotypic changes at multiple target loci [1]. If regulatory GP maps evolve under this type of selection, what kind of resulting maps should we expect to observe?

Many broad questions about gene regulatory architecture remain unanswered and might benefit from a suitable evolutionary framework. It is not clear why TFs have such short motifs in eukaryotes [2] and rely on intricate mechanisms for promoter search [3, 4], why so much of human DNA is transcribed [5], or why regulatory networks are so densely interconnected by weak links [6]. A number of recent studies that characterize the genotype-expression maps of promoters or enhancers in *E. coli* [7, 8], *S. cerevisiae* [9, 10], and *D. melanogaster* [11, 12] found that a large fraction of random DNA sequences can act as a promoter or an enhancer, and even those that do not are only a few point mutations away. Should we be surprised by this finding?

In this paper, we study the evolution of “regulatory parameters,” i.e., parameters that determine the gene regulatory architecture shared by all regulatory sequences. For example, these could be parameters associated with TFs (such as their concentration, binding site length, or preferred binding motif sequence), which affect the GP map of multiple regulatory sequences across the entire genome. Alternatively, these could be parameters determining the major architectural features of the regulatory apparatus, such as the typical number of TF binding sites in a CRE required to successfully control gene expression [13], or the presence/absence of mechanisms that allow broad and reliable silencing of large gene clusters [14]. In detail, we will focus on how the space of possible DNA sequences at a target locus is divided among the relevant gene expression phenotypes; for example, what fraction of possible CRE sequences binds or doesn’t bind a particular TF?

The essential premise of our theory asserts that different CREs should not be viewed as evolving on independent “fitness landscapes,” although this would be the standard approach for protein structure or TF-DNA binding landscapes [15, 16]. While selection does act differently on individual CREs — each CRE needs to express genes in a condition- and tissue-specific manner —, the map from CRE sequence to expression given TF concentrations should be treated as a single GP map. This is because it arises from the shared molecular machinery operating under universal biophysical laws that govern protein-DNA interactions: colloquially, it is precisely the structure of these interactions that we refer to when we discuss “genetic architecture”. Quantitatively, such architecture is defined by a common set of regulatory parameters *λ*, which are also encoded at various “regulatory loci” somewhere in the genome, and can therefore evolve (Fig. 1). In other words, all CREs share the same genotype-to-phenotype map conditional on regulatory parameters and TF concentrations, but might have individual selection (fitness) requirements.

**Figure 1:**
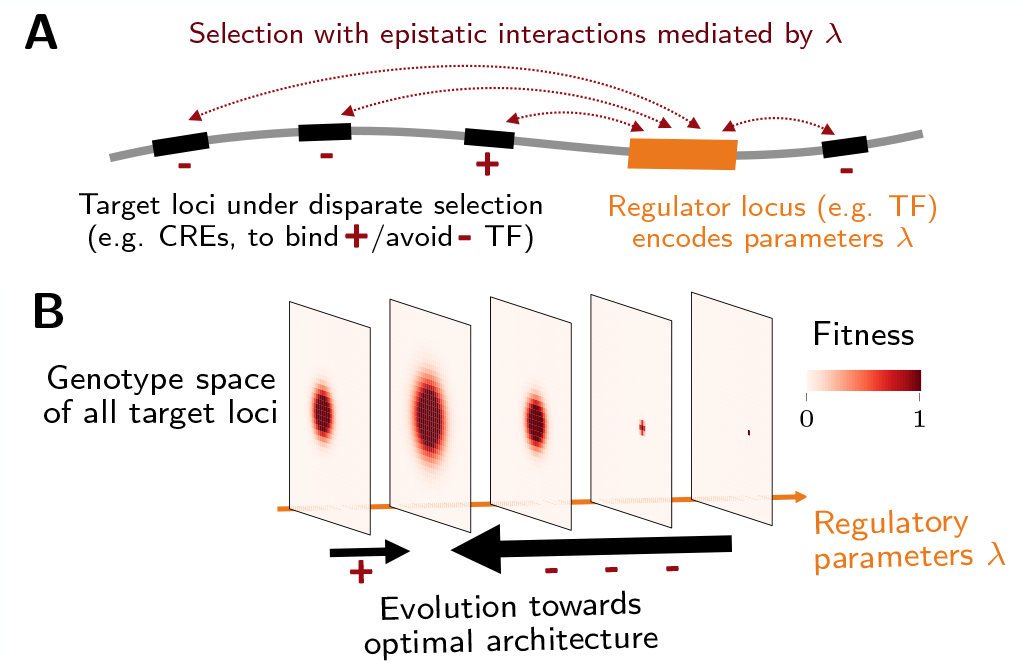
A summary of the main argument. **(A) Global view.** Multiple regulatory sequences such as CREs (“target loci,” black) are under selection for diverse phenotypes, e.g., to bind or avoid a TF (red plus or minus). The gene expression phenotypes at different CREs are influenced by “regulatory parameters” *λ*, which are encoded at “regulator loci” elsewhere in the genome, e.g., in a TF gene (orange). As a result, the regulator and the target loci co-evolve, with epistatic interactions via *λ* (dotted arrows), on a fixed global landscape, jointly describing all target and regulatory loci. **(B) Local view**. In this alternative view, we can look at a marginal genotype-phenotype map at a single target (CRE) locus, keeping in mind that the same GP map is replicated across all the CREs in the genome using shared regulatory parameters. We argue that the regulatory parameters *λ* (horizontal axis) that determine this replicated GP map will evolve towards an optimum, where a large fraction of genotypes at the target loci (vertical planes) have high fitness (red shading). This “optimal architecture” will reflect the statistics of phenotypes selected for at different target loci, e.g., by finding a compromise between sites under selection for and against TF binding. Even on such optimal architecture, selection must still act at each individual CRE, to adapt its sequence to its particular functional requirement. Note that while this schematic implies that fit genotypes cluster together in a single fitness peak, that is not required by the theory in the weak mutation limit; optimal maps may feature multiple degenerate peaks.

If we include the regulatory loci into our consideration, we can further imagine a single, global, unchanging GP map that contains all CREs plus the regulatory loci. For our theoretical approach, such a global, fixed landscape view is indispensable, because it allows us to formally separate, or jointly treat, at will, the coevolution of individual CREs (Fig. 1A, black) and of the shared regulatory parameters (orange). For the readers’ understanding, it is essential to also navigate the complementary, “local” view, more suitable for data analysis and simulation of a single CRE, which we adopt in the companion paper [17]. In the “global” view, a single GP map for all CREs and, simultaneously, for the regulatory parameters, is taken as given while the adaptive dynamics unfolds on it. In the “local” view (Fig. 1B), the marginal, effective GP map and fitness landscape for an individual CRE changes as the regulatory parameters evolve; yet one must constantly think of the same GP map as being “replicated” self-consistently for each CRE across the genome. We will refer to this structure, where the shared regulatory parameters *λ* shape a single GP map replicated across multiple loci (here, CREs) in the genome, each of which can be subject to different selective pressures, as a “replicated genotype-phenotype map.”

Our key finding can be summarized in a simple principle: on replicated GP maps, evolved regulatory parameters associate fit phenotypes with a large number of genotypes (Fig. 1B). This provides a bridge between population genetics and optimality theories – it is a way to derive optimal parameter values rooted in population genetics theory. We will see that, surprisingly, selection can drive regulatory parameters towards their optima even outside the classical limit where selection on each CRE locus is much stronger than drift.

Our approach builds on an analogy between population genetics and statistical physics [18, 19], in particular the theory of free fitness developed by Iwasa [20] and Sella and Hirsh [18], who in turn built on Wright’s equilibrium distributions [21] and Kimura’s fixation probabilities [22]. We furthermore draw strongly on our recently established links between evolution and information theory [23]. Free fitness is a quantity that is maximized as evolution approaches equilibrium. We use this result to express a trade-off between fitness and genetic information, a measure of evolutionary constraint on the genotype (the difference between the genotype distribution under selection vs. under neutrality). For a given genotype-phenotype-fitness map, there is a minimal information required for any given expected fitness, with the population size controlling the tradeoff between information and fitness. Regulatory parameters that determine the GP map are under selection to maximize free fitness.

Conceptually, our results lead to two levels of optimization, which emerge automatically within a single evolutionary framework. Regulatory parameters are optimized for “free fitness” and will shape the GP map so that its statistical properties match what selection might favor, in aggregate, across all CREs in an organism. Given those parameters, selection will further act, individually, on every CRE, to adapt its unique sequence to its particular functional requirement that will contribute towards maximizing organismal fitness. If the GP map is optimized as a whole via regulatory parameters, selection needs to exert least “work” to fix and maintain genotypes at individual CREs in order to implement their desired function.

We develop our theory and illustrate it in parallel on a toy model of gene regulation, which we refer to as the “simple binary model of TF binding”, in Section 1. Starting from basic population genetics assumptions, we derive an optimization principle for the regulatory parameters of the replicated GP map, and connect it to optimal coding in information theory. The assumptions are then questioned in Section 2, where we discuss how biophysical constraints on gene regulation affect the information requirements, and how the presence of strong mutation adds robustness as an additional factor in the optimization. In this section, we illustrate the biologically relevant complications by extending our simple binary model from Section 1 in various directions.

## 1 Theory for optimal genotype-phenotype maps

The theory contains three key ingredients that we consider below. First, a class of population genetics models that is very general but still tractable at stationarity. Second, an analysis analogous to energy-entropy trade-off in statistical physics. And third, an interpretation of this trade-off as an optimization principle relevant for evolution on replicated GP maps, as is the case for genetic regulatory sequences. Finally, we point out a connection to optimal coding in information theory, which provides an intuition about what the optimal GP maps for regulation should look like.

### Population genetics setting

We track a haploid population with effective size *N* that evolves under selection, mutation and drift. Initially, we work in “the fixed states approximation”, which requires a low enough mutation rate, *NLu* ≪ 1, where *L* is the size of genetic regulatory sequences considered, and *u* is the mutation rate (per bp, per individual, per generation); we relax this assumption later.

In the fixed states approximation, the population state is given by the most recently fixed genotype (labeled *i, j*, …), and evolutionary stochasticity is captured by the distribution over these genotypes, *ψ*_*i*_. Mutations are proposed at a rate *u*_*ij*_ (from *i* to *j*), and accepted with Kimura’s fixation probability [18, 22] that depends on the log-fitness values *w*_*i*_ and *w*_*j*_ of the respective genotypes (also referred to as “additive fitness” in Ref. [18]).

The resulting dynamics is a continuous time discrete space Markov chain, and we are interested in its equilibrium distribution, *ψ*_*i*_. For this we assume that *N* and *w*_*i*_ are constant over time, and that the mutation rates *u*_*ij*_ satisfy detailed balance. Then, the probability *ψ*_*i*_ of being fixed for the genotype *i* has a simple form [18, 19, 21, 24]:

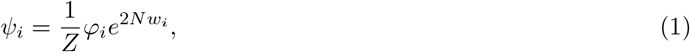

where 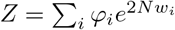 is a normalization constant and *φ*_*i*_ is the equilibrium distribution under neutrality (i.e., if all fitness values *w*_*i*_ were equal). Under the simplest model of only single nucleotide replacements and no mutation bias, *φ*_*i*_ is uniform over all sequences of a given length. Often, we can simplify the description by grouping all genotypes by their log-fitness: *w* occurs with probability 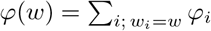 under neutrality and *ψ*(*w*) = *φ*(*w*)*e*^2*Nw*^*/Z* under selection.

Taken together, systems at equilibrium are fully specified by the fitness landscape *w*_*i*_, the neutral distribution *φ*_*i*_, and the population size *N*. Selection increases the probability for genotypes with high fitness, and is more efficient in larger populations. This is analogous to the Boltzmann distribution in physics, with log-fitness instead of negative energy and 2*N* instead of inverse temperature.

In Fig. 2A-C, we illustrate the key aspects of equilibrium behavior in a simple (yet, surprisingly rich) binary model for TF binding. The model consists of *n* CREs, *nf* of which are under selection to bind a TF, while the remaining *n*(1− *f*) are under selection to avoid it. TF binding is treated as a binary phenotype, and each CRE with the wrong phenotype (FN, a false negative, for each failure of a TF to bind where required; FP, a false positive, for each ectopic TF binding where unwanted) incurs an equal log-fitness penalty *s* per wrong phenotype at a CRE; a more refined model could easily introduce asymmetric penalties for FNs and FPs. As a result, the log-fitness is 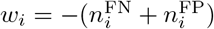*s*, where 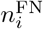 and 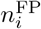 are the counts of both kinds of error in genotype *i* (see Fig. 2A). We assume that all CREs share the same replicated GP map, and that under neutrality, a TF would bind a fraction *q* of all possible CRE sequences (Fig. 2B). This yields the neutral distribution of log-fitness, *φ*(*w*), since errors will be binomially distributed, 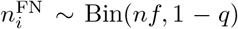 and 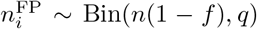. Under selection, the distribution *ψ*(*w*) is pushed towards higher *w*, depending on the population size (Fig. 2C).

**Figure 2:**
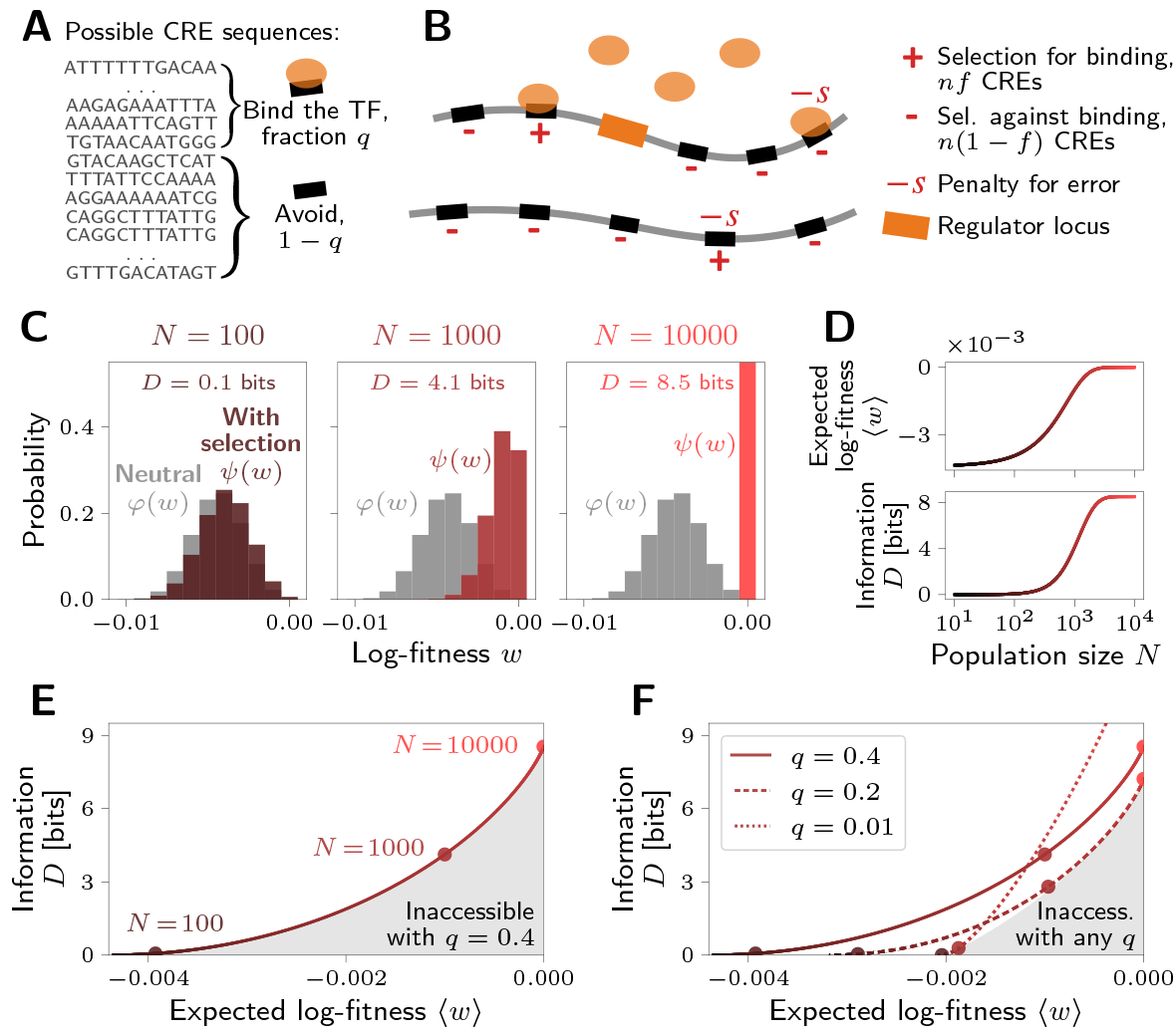
A simple binary model of TF binding on a replicated GP map. **(A)** The entire genotypic space for CREs contains 4^*L*^ possible sequences of length *L*, among which a fraction *q* binds and 1 − *q* avoids a given TF. *q* is a regulatory parameter. **(B)** The concrete organismal genotype contains *n* CREs (black sequence regions, here *n* = 10), of which a fraction *f* (here *f* = 0.2) is under positive selection to bind the TF (orange plus) and 1 − *f* under negative selection, favoring TF not being bound (orange minus). Each error (“false positive” and “false negative”, see main text) reduces log-fitness by the same amount, *s* (here, *n*^FP^ = *n*^FN^ = 1, implying *w* = − 2*s*). The single regulatory parameter *q* of this simple binary model is encoded at a regulator locus (orange; e.g., the TF gene that affects its concentration). **(C)** Distributions of log-fitness under neutrality (gray, *φ*(*w*)) and under selection (red, *ψ*(*w*)) at different effective population sizes *N* (shade of red, increasing left-to-right). Parameters as in (B), with *s* = 0.001 and *q* = 0.4. **(D)** Expected log-fitness ⟨*w*⟩ (top) and information *D* (bottom) both increase with increasing population size. **(E)** A parametric curve of the expected log-fitness ⟨*w*⟩ and information *D* with increasing population size *N* (black-red gradient) for a fixed *q* = 0.4. Equilibrium distributions have the lowest possible *D* at a given ⟨*w*⟩ and define the boundary. Non-equilibrium distributions above the boundary exist but require extra information for the same average log-fitness; the region below the boundary is inaccessible to any realistic process at fixed *q*. **(F)** If the marginal genotype-phenotype map for a CRE changes (via a change in the regulatory parameter *q*), new combinations of ⟨*w*⟩ and *D* are unlocked compared to fixed *q* = 0.4 (gray region shrinks compared to E). Nevertheless, a certain region of the fitness vs. information plane still remains inaccessible for every *q* (gray). Dots on the boundaries correspond to *N* = 10, 1000, 10000, as indicated in E.

In summary, our model is the simplest example of a replicated GP map that applies simultaneously across *n* CREs in the genome, even though the selection acting on each CRE may be different – some are required to bind a TF while others are required not to bind it. The replicated GP map is parametrized by a single regulatory parameter, *q*. The subsequent assignment of fitnesses to phenotypes at each CRE requires two additional parameters, *s* and *f*. This model is intentionally simple for illustration purposes; later we introduce continuously varying binding probabilities, multiple TFs, and non-CRE loci. Note, however, that our analysis does not depend on exactly which CRE sequences bind the TF, or on the physics of TF binding itself, in contrast to previous studies that employed similar evolutionary models in more mechanistic settings [1, 18, 24–26]. In our setup, details such as the CRE length, TF binding motif or possible cooperativity between binding sites are all subsumed into a single regulatory parameter of the GP map, *q*.

### Fitness-information trade-off

As the population size *N* increases, selection becomes more efficient, and the population achieves a higher expected log-fitness ⟨*w*⟩ =∑ _*i*_ *ψ*_*i*_*w*_*i*_. To this end, the distribution *ψ*_*i*_ must become increasingly concentrated at genotypes with high fitness. We quantify this as “genetic information” *D* – a Kullback-Leibler (KL) divergence [23, 27]

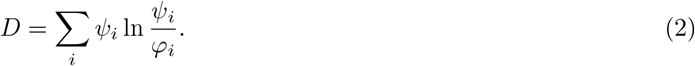

*D* quantifies how the distribution under selection, *ψ*, differs from that under neutrality, *φ*, and generalizes several previously proposed measures [2, 28, 29]. If *ψ* and *φ* are equal, e.g., in absence of selection or before it had time to act, *D* is zero. Whenever *ψ* and *φ* differ, *D* is positive. In the important special case of a uniform neutral distribution *φ*_*i*_, *D* is equal to the reduction in Shannon entropy, *D* = *H*(*φ*) − *H*(*ψ*). Intuitively, higher *D* means that evolution is constrained to a smaller fraction of the genotype space and selection has to do more “work” to find and fix high fitness genotypes.

Conveniently, the information *D* can be computed or lower bounded using distributions over phenotypes or fitness [23], which is what we do in Fig. 2C. Accumulated information *D* and average log-fitness ⟨*w*⟩ both increase with population size *N* (Fig. 2D), and trace out a parametric curve in the fitness-information plane (Fig. 2E).

Crucially, the region below this curve is inaccessible at any fixed value of the regulatory parameter *q* of the GP map. This can be derived rigorously, by showing that the equilibrium distribution in 1 maximizes free fitness [18, 20, 23], which can be conveniently expressed in dimensionless units (or, equivalently, converted into bits) in terms of mean log-fitness and genetic information:

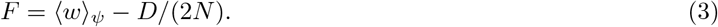

In summary, evolution tends to achieve any given mean log-fitness ⟨*w*⟩ with the least sequence constraint (information) possible; or conversely, the highest possible log-fitness at any given information *D*. The relevant mathematics is strongly related to the maximum entropy principle [30–34].

### Regulatory parameters evolve towards optimal values

The equilibrium theory implies a minimal *D* for any ⟨*w*⟩ in a given system, but this minimal value will change if the GP map itself changes. In our simple binary model, the only regulatory parameter that shapes the GP map is *q*, the fraction of sequences within the entire genotype space that bind the TF. We plot the fitness-information trade-off curve for several different values of *q* in Fig. 2F. Clearly, the inaccessible region at any fixed *q* can be shrunk if *q* itself is allowed to vary. Such variation is biologically plausible: for example, *q* could evolve via changes to the TF’s own regulatory sequence (making the TF concentration higher or lower, causing the TF to generically bind more or less sequences), or to the TF’s DNA-binding domain (making the TF bind more or less promiscuously at a fixed concentration). As a whole, the entire family of marginal GP maps parameterized by the regulatory parameter *q* can access higher average log-fitness values at a given information *D* than any single map at a fixed *q* value. Yet even with the ability to change *q*, certain values of ⟨*w*⟩ will remain out of reach. What can we say about evolution of the regulatory parameter *q* in our model?

Fig. 2F suggests that some values of *q* are more advantageous than others, depending on the population size *N*. For example, in small populations, selection within CREs is ineffective, marked by *D* close to zero. It seems best to choose *q* such that even random sequences (“random CREs”) have higher expected fitness ⟨*w* ⟩. In the simple binary model, this means choosing a vanishingly small *q*, because the majority of CREs are under selection against binding (*f <* 0.5). Large populations, in contrast, have the capacity to evolve the desired binding phenotype at each CRE individually. In this case, ⟨*w*⟩ is anyway near maximum. It seems best to choose an intermediate *q*, such that all the desired phenotypes are well represented in the CRE genotype space, and adaptation requires as little information *D* as possible. Intuitively, therefore, adjusting the regulatory parameter *q* can adjust the distribution of phenotype values across all CREs as close as possible to the selected phenotype distribution across all CREs, but only at the aggregate, statistical level, not CRE-by-CRE. At low *N*, this is the best that can be done. At high *N*, selection can further reshape this overall distribution to match each individual CRE precisely to its desired phenotype; if that is possible, a different *q* at high *N* will be preferred than at low *N*. This reasoning can be generalized to claim that regulatory parameters such as *q* have an optimal value, as a function of population size, which maximizes free fitness (Eq. 3). A union of such optima for different *N* comprises the boundary between the accessible and inaccessible regions in Fig. 2F, and we show their values in Fig. 3A.

**Figure 3:**
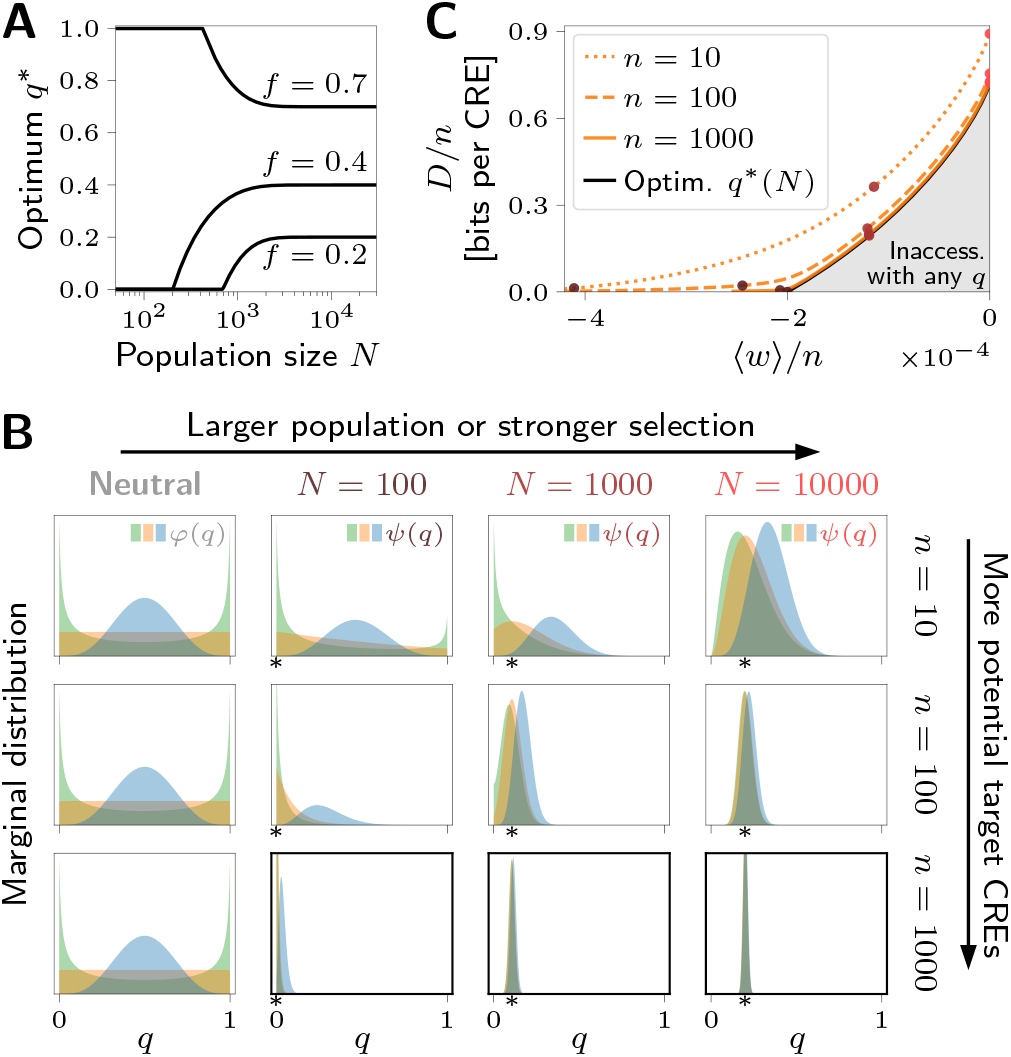
Regulatory parameters of replicated GP maps can be strongly optimized even at weak selection on individual target loci. **(A)** Optimal values for the regulatory parameter *q* as a function of population size *N* maximize free fitness in the simple binary model of Fig. 2. Optimal values depend on the fraction of CREs under selection to bind the TF (here shown for *f* = 0.2, 0.4, 0.7). At high *N*, we generically observe “frequency matching” solutions, where *q*^*\**^ = *f*. **(B)** Routes towards optimality for the regulatory parameter *q*. Three example neutral distributions *φ*(*q*) are shown in the left column (colors). Under selection, equilibrium distributions *ψ*(*q*) depend on population size *N* (left to right) and the number of target CREs *n* (top to bottom). With increasing *N, ψ*(*q*) universally peak, as expected, at the optimum value *q*^*\**^(*N*) (asterisks on horizontal axes), as in (A). Alternatively, strong optimization with tight regulatory parameter concentration around the maximum also happens at modest *Ns*, if many target loci *n* are present (bottom row). **(C)** As the number of target loci (CREs) *n* on the replicated GP map increases (orange curves) and the regulatory parameter *q* is driven towards its optimal value *q*^*\**^(*N*) for each *N* as shown in (A), the information vs mean log-fitness curves, properly rescaled by *n*, converge to the optimal information/fitness tradeoff (black). Orange dots highlight (*N, n*) combinations depicted in (B). Fixed parameters: *f* = 0.2, per-target-locus selection coefficient *s* = 0.001.

But does evolution actually drive regulatory parameters towards these optimal values? To answer this, we need to take a global view and include the loci controlling the regulatory parameters into the theory. We will refer to these loci as “regulator loci” and label their genotype with *r*, while the remaining “target loci” keep the label *i*. Their joint equilibrium distribution is 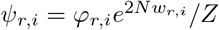. We assume that the two sets of loci do not overlap, such that under neutrality, *r* and *i* are independent and *φ*_*r,i*_ = *φ*_*r*_*φ*_*i*_. We are especially interested in the evolution of the regulatory loci. The marginal distribution over *r* is

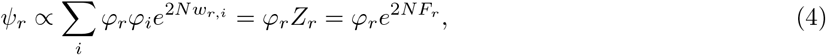

where 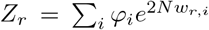 is the partition function for the target loci conditional on a regulatory genotype *r*, and *F*_*r*_ = ln *Z*_*r*_*/*(2*N*) = ⟨ *w*_*r*_⟩ −*D*_*r*_*/*(2*N*) is the corresponding free fitness. Therefore, while the equilibrium distribution of the entire system analyzed in our “global view” has log-fitness of entire genotypes in the exponent as in 1, the marginal distribution of regulatory loci that interact with target loci has (conditional) free fitness in the exponent. Intuitively, advantage goes to regulator genotypes that achieve high fitness in combination with a large number of target genotypes *i*, and the interplay of fitness and numbers (analogous to the energy–entropy balance in statistical physics) is captured by free fitness [35].

The interaction between the regulatory loci and their targets is often mediated only via a collection of regulatory parameters *λ*, and we might not know (or care) how these are encoded in the genome. (Our simple binary model has only a single regulatory parameter, *λ* = {*q*}, and we need not commit to any molecular mechanism about how *q* might be encoded in the genome). In that case, we can write log-fitness in 4 as *w*_*i*_(*λ*), conditional free fitness as *F* (*λ*), and summarize the possibly complicated genetic architecture of *λ* as a neutral density of states, *φ*(*λ*) = ∑_*r*; *λ*(*r*)=*λ*_ *φ*_*r*_. This expresses how common different values of *λ* would be under neutrality, i.e., among random DNA sequences. Under selection, the equilibrium distribution of regulatory parameters *λ* will be:

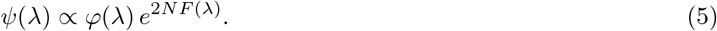

5 is similar to, yet subtly and importantly different from, the classic equilibrium results that follow directly from 1. In those setups (e.g., Refs. [24, 25]), the neutral density of states can be overwhelmed as population size *N* or selection strength increase, driving the evolution towards fit solutions. Since free fitness replaces log-fitness in 5, new phenomenology emerges when *Ns* remains constant but free fitness nevertheless grows as the number of regulator targets, *n*, increases—this is because *F*_*r*_ is a sum over all CREs and thus scales linearly with *n*. Therefore, not only in large populations, but also in finite populations with many target loci governed by a single replicated GP map, the factor *e*^2*NF* (*λ*)^ will overwhelm the neutral density of states and *λ* will be strongly optimized for the free fitness of its targets.

This new route towards optimality is illustrated for our simple binary model in Fig. 3B. On the left are shown three examples of a possible neutral distribution *φ*(*q*), chosen arbitrarily. Towards the right, we increase the population size and observe how selection on the target CREs drives the distribution *ψ*(*q*) towards a form increasingly independent of the neutral distribution, and dominated by free fitness. As reported in Fig. 3A, the optimal *q* depends on *N*→ ∞. Beyond around *N* = 10000, further increases in *N* would not make a difference: all CREs are adapted to their required phenotype, and the distribution *ψ*(*q*) is dominated simply by what fraction of possible genotypes realize the required combination of phenotypes. This *Ns* limit, where selection on each CRE locus is stronger than drift, is the classic route towards evolutionary optimization. An alternative route is shown at fixed – even modest – *Ns*, as the number of target CREs, *n*, increases, indicated in Fig. 3B by going top-to-bottom. Because the conditional free fitness *F* (*λ*) is extensive in the number of target CREs, not only is the neutral distribution *φ*(*q*) overwhelmed as *n* grows, such that *ψ*(*q*) increasingly centers around a clear optimum *q*^*\**^(*N*); *ψ* furthermore becomes increasingly peaked around the optimal value *q*^*\**^. Therefore, unless physically impossible, we would expect that regulatory parameters of replicated GP maps that govern many target loci are very strongly optimized at equilibrium. In other words, while the optimal regulatory parameter value can depend on *N*, the evolutionary drive towards this optimum can happen also when selection per locus is weak relative to drift, so long as *n* grows large, so as to optimally balance the information vs mean log-fitness tradeoff of Fig. 3C.

### Optimal genotype-phenotype maps and analogies to coding

If GP maps are indeed optimized, what form will these maps have? At the optimal choice of regulatory parameters, how much information would selection subsequently still need to accumulate in the target loci to reach the fittest solutions?

To address these questions and develop intuition for the general case where multiple discrete phenotypes are possible, we now introduce the first extension of the simple binary model. We consider multiple discrete phenotypes *z* ∈ {0, *Z*} (of which the simple binary model was a special case, with *Z* = 1). One phenotype is specified by each of the *n* target loci, such as CREs or even non-CRE loci, which share the same replicated GP map. Across these loci, a fraction *q*_*z*_ realize the phenotype *z* under neutrality (i.e., a phenotype *z* is specified by a fraction *q*_*z*_ of all sequences). *Z* + 1 regulatory parameters *λ* = {*q*_*z*_} are all allowed to vary to maximize free fitness, up to the constraint 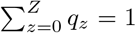, such that phenotypes cannot overlap in sequence space. Of the *n* target loci, fractions *f*_*z*_ are forced by strong selection to evolve the phenotype *z*.

When selection per target locus is stronger than drift (large *Ns* limit), maximization of free fitness, *F* (*λ*) = ⟨*w*(*λ*) ⟩− *D*(*λ*)*/*(2*N*), implies minimization of *D*(*λ*), since the expected log-fitness ⟨*w*(*λ*) ⟩ is anyway at its maximum. How should the genotype space be divided among the phenotypes, so that the required phenotypes are realized by the largest number of genotypes, i.e., with the lowest *D*(*λ* = {*q*_*z*_})? Given *q* and *f*, the information is *D*({*q*_*z*_}) =*∑*_*z*_ *nf*_*z*_ ln(1*/q*_*z*_), since *nf*_*z*_ target loci must realize *z* with probability 1, while under neutrality only a fraction *q*_*z*_ of target loci will do so. The optimal partitions 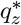 are given by

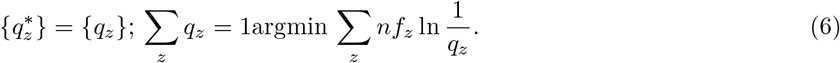

This minimization problem is identical to the classical problem of optimal codes in information theory, dating back to Shannon [36]. A source generates messages *z* with probabilities *f*_*z*_, and each is assigned a code word that is *l*_*z*_ symbols long. We want to choose code words such that the average length, ∑_*z*_ *f*_*z*_*l*_*z*_ is minimized. In6, ln(1*/q*_*z*_) corresponds to the lengths *l*_*z*_, and the condition ∑_*z*_ *q*_*z*_ = 1 corresponds to the Kraft inequality [27], a requirement for unique decodability of the original messages. (Kraft inequality is _*z*_ *q*_*z*_ ≤ 1 but equality is achieved at the optimum; also, in information theory, *e* and the natural ln are usually replaced by 2 and log_2_ to express code length in bits). The solution is

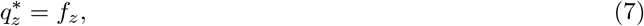

with minimal information

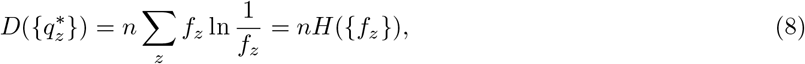

where *H*({*f*_*z*_}) is the Shannon entropy of probabilities *f*_*z*_. Therefore, on an optimal replicated map under strong selection, the fraction of all sequences that encode each phenotype should match the desired frequency of that phenotype. This statement generalizes and makes precise the “frequency matching” nomenclature we used for optimal solutions at high *Ns* in Fig. 3A. Furthermore, on an optimal map, the minimal amount of genetic information needed is given by the entropy of desired phenotype frequencies. Information theory asserts that this is an absolute bound. In the coding analogy, each target locus must encode a “message” about which phenotype should be realized there, and the minimal amount of information required is proportional to the optimal code length needed to transmit such messages. In engineered systems, information theory is used to devise practical codes that approach this optimal bound [27]; apparently, strong selection is similarly driving organisms to efficiently solve an analogous coding problem, by evolving novel regulatory mechanisms that approach a mathematically identical bound. Architectures reaching such a bound would be “IT-efficient” and thus the best possible, in an absolute sense.

On an optimal GP map, rare messages, i.e., phenotypes such as TF binding that is required only in a minority of target loci, require a considerable amount of information to encode, ln(1*/f*_*z*_) *>* 1 per target locus (Fig. 4A). In contrast, phenotypes that are common (e.g., “TF should not bind here”) take little information, *f*_*z*_ ≈ 1 ln(1/*f*_*z*_) ≈ 0. Even across the entire genome, the most common phenotype requires less information to encode than the rare phenotype (Fig. 4B). However, it is precisely the need to encode the common phenotype cheaply that will have shaped the optimization of the entire replicated GP map towards devoting most sequence space to them, leaving less space for the rare but nonetheless important (and costly, information-wise) phenotypes.

**Figure 4:**
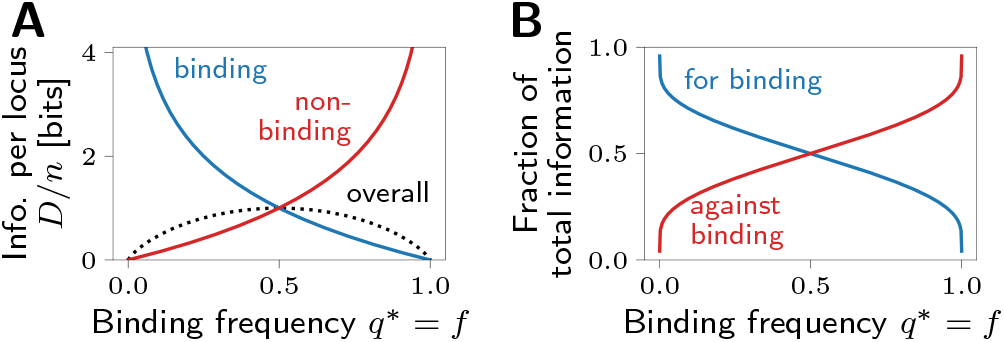
Information needed to fulfill the required state for or against TF binding at each CRE target locus. Results are shown for the simple binary model of Fig. 2, assuming the optimal “frequency matching” (*q*^*\**^ = *f*) regime at high *Ns*. **(A)** Information per CRE under positive (blue) and negative (red) selection, and the overall average per CRE (black dotted line). Phenotypes that are rare (e.g., binding when *f <* 0.5) require more information per CRE than phenotypes that are more common. **(B)** Encoding the rarer phenotype (e.g., TF binding when *f <* 0.5) requires more information also in total, despite the smaller number of loci where it occurs.

## 2 Towards optimal maps with realistic mechanisms

The idealized scenario that led us to the optimal coding analogy in the large *Ns* limit, as well as the population genetics assumptions needed to derive our free-fitness-based theory, depend on strong simplifications. Here we show how the theory can be extended to cover three important aspects of real systems. **First**, if the phenotype *z* is highly multidimensional (e.g., if *z* is the binding status of many TFs at a target locus), biological mechanisms may be insufficiently flexible to independently tune all the probabilities *q*_*z*_ and achieve the IT-efficient optimum of Eqs. (7,8). Nonetheless, we will argue that novel regulatory mechanisms could evolve to push populations much closer towards, even if not strictly to, the information-theoretic optima. **Second**, phenotypes are often not discrete, but quantitative or probabilistic (e.g., TF molecules bind DNA with some sequence- and parameter-dependent probability). We show that this can lead to “overspecification”: extra information will need to be accumulated and maintained by selection to guarantee the desired phenotypic outcome. **Third**, we question the assumption of low mutation rate that underpins the fixed states approximation. We outline how our results change in genetically diverse populations, but postpone a full theoretical development for the future.

### Availability of regulatory mechanisms

The IT-efficient frequency-matching solution of 7 may be biologically unachievable because the available regulatory mechanisms do not have sufficient flexibility to individually adjust the probability *q*_*z*_ of each phenotype *z*. We explore this by extending our simple binary model of TF binding yet again (Fig. 5A), as follows. First, we consider a larger number, *T >* 1, of TF types. A particular TF must bind some fraction *f* of the CRE loci, and avoid the rest of the CRE loci (subject to a penalty *s* per error), as before. Since there are *T* TF types, CRE loci are under selection to be bound by *f T* TFs. Second, we introduce a set of *m* non-CRE target loci into the model. These new non-CRE loci share the same replicated GP map as CRE loci, but are under selection to avoid any TF binding whatsoever (for simplicity, also under penalty *s*). Under neutrality, any single TF binds to any (CRE or non-CRE) target locus with probability *q*, independently of other TFs.

**Figure 5:**
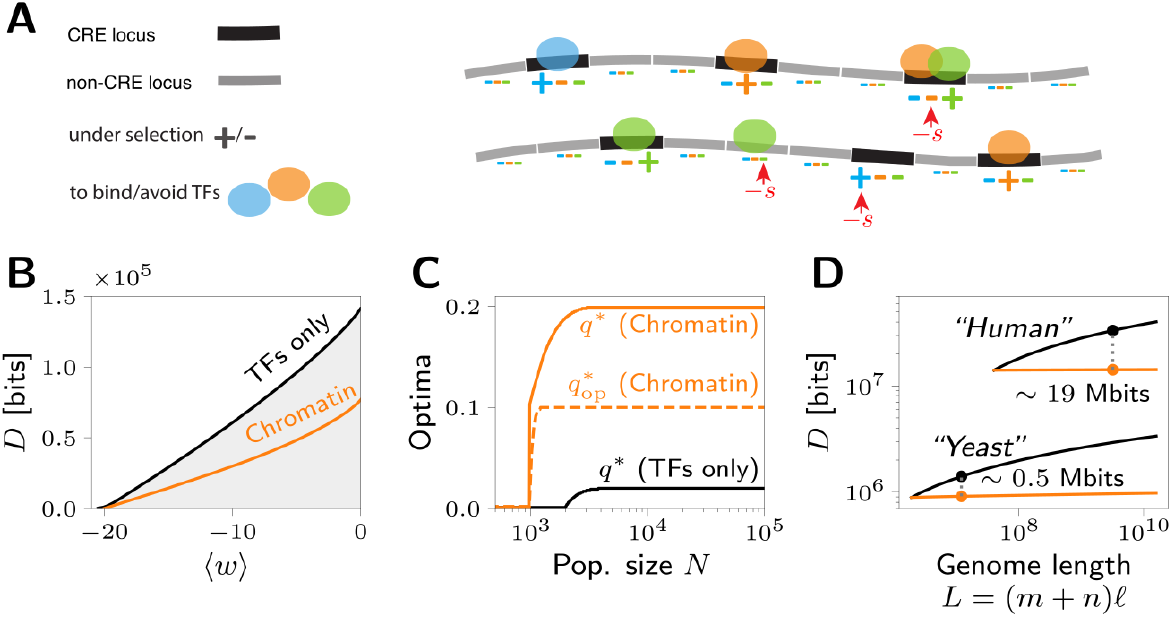
Novel regulatory mechanisms can improve coding efficiency. **(A)** An extended model with *T* TFs, *n* CREs and *m* non-CRE loci (here, *T* = 3, *n* = 6 and *m* = 14). Each CRE is under positive selection for a fraction *f* of TFs and negative selection for 1 − *f* (here *f* = 1*/*3), while non-CRE loci are under negative selection only. Spurious binding (“false positive,” FP) and failure to bind where required (“false negative,” FN) both reduce log-fitness by *s* (red), as in Fig. 2. **(B)** Fitness-information trade-off as in Fig. 2F, shown without (black) and with (orange) the additional layer of chromatin regulation. Values of *q* and *q*_op_ are optimized for each population size and shown in (C). Chromatin considerably reduces the information required for any given fitness. Parameter values are *n* = 1000, *m* = 9000, *T* = 100, *f* = 0.1, *s* = 10^*−*3^. **(C)** Parameter optima for the models used in (B). **(D)** Comparison of information needed at strong selection, with and without chromatin. Parameter values are inspired by the human and yeast genome (see text). Savings due to chromatin are highlighted by vertical dotted lines.

Note that there are alternative biological setups and verbal descriptions that would distill into the same mathematics. Instead of individual TF binding profiles at target loci, one could think of these same loci being transcribed under different “conditions”. Similarly, our arguments are not dependent on narrow molecular interpretations of “binding” or “not binding”, and could be extended to cooperative interactions, multiple TF binding sites for the same TF etc. For clarity of theoretical exposition, we do not pursue these generalizations here.

Under neutrality, the probability for a CRE to bind the correct set of *f T* TFs and avoid the other (1 − *f*)*T* TFs is *q*^*fT*^ (1 − *q*)^(1*−f*)*T*^. A non-CRE locus successfully avoids all TFs with probability (1 − *q*)^*T*^. For a given value of *q*, we can again compute ⟨*w*⟩ and *D* as a function of population size (Fig. 5B, black).

What GP map will evolve for this extended model? At low *N* and with *f <* 0.5, *q*^*\**^ is zero, as before (Fig. 5C, black). At high *N*, the ideal solution is frequency matching. Perfect frequency matching divides the sequence space among (at most) *n* + 1 required phenotypes: *n* binding profiles for *T* TFs (at most one unique profile per CRE) and 1 “avoid all” phenotype for all *m* non-CRE loci. But it is certainly impossible to generically realize such frequency matching by varying a single parameter *q*. Even hypothetically considering more complex yet still realistic models of gene regulation, we (or selection!) might not be able to single out the required sets of TF combinations, out of the 2^*T*^ possible ones, to generate an optimal GP map by adjusting the regulatory parameters alone. In other words, in this extended model, biophysical constraints prevent selection from accessing an IT-efficient frequency matching solution for the replicated GP map. As a consequence, selection will need to do additional work at each individual target locus to adapt it, on a suboptimal GP map, at a much higher total information cost *D*.

In our model, varying *q* can only lead to more or less binding overall. Under free fitness minimization, *q* will evolve to *q*^*\**^ = *nf/*(*n* + *m*), matching the fraction of all target loci that need to bind any particular TF, and the necessary information will be

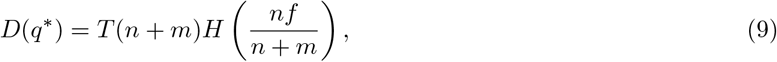

proportional to the number of TFs. This arrangement is very inefficient from the perspective of the non-CRE loci, which must avoid each TF separately. This does not necessarily mean that the total information in non-CRE loci is high. Instead, the need to avoid them all means that the optimal *q*^*\**^ is low, leading to high information requirements for binding TFs inside CREs. For example, with *q*^*\**^ = 0.01, about 82% of *D*(*q*^*\**^) is inside CREs (87% with *q*^*\**^ = 0.001 or 71% with *q*^*\**^ = 0.1; using order-of-magnitude parameter values reported in Fig. 5 caption).

Can organisms reduce the required information by introducing an additional layer of regulation? For our extended model, such an extra mechanism could implement a switch deciding between two states: either leaving a CRE permissive to TF binding, or vetoing any and all TF binding at a non-CRE locus in a single regulatory step [14]. For simplicity, we refer to these two states as open and closed chromatin. We realize that other molecular scenarios could be invoked (methylation and CpG islands [37], the histone code, etc.), or could contribute towards regulating the chromatin state [37–39].

With the new mechanism, the replicated GP map gains one more regulatory parameter *q*_op_, so that *λ* = {*q, q*_op_}. Under neutrality, there is a probability *q*_op_ for a target locus to be open and 1 − *q*_op_ to be closed. Therefore, CREs achieve their required phenotype with the probability *q*_op_*q*^*fT*^ (1− *q*)^(1*−f*)*T*^. Non-CRE loci achieve their required phenotype with probability 1− *q*_op_ + *q*_op_(1− *q*)^*T*^ ≈ 1− *q*_op_ – either by being closed, or by being open but avoiding all TFs; but the latter option has negligible probability with an appreciable number of TFs. This leads to a considerable improvement in the fitness-information trade-off (Fig. 5B, orange). In the frequency-matching regime, the optimal values are 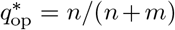, matching the frequency of CREs that need to be open to binding, and *q*^*\**^ = *f*, matching the fraction of CREs that any TF needs to bind (Fig. 5C, orange). The necessary information is

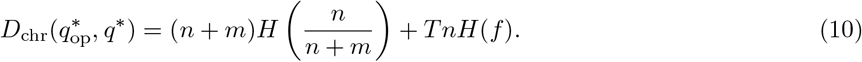

The first term is the information needed to specify which target loci are CREs; importantly, this does not grow with the number of TFs. The second term is the information needed to specify which CREs must be bound by each of the *T* TFs; this is independent of the number of non-CRE loci.

The addition of a chromatin-like mechanism can lead to a dramatic reduction in the required information compared to the single-layer regulation in 9. We illustrate the difference in Fig. 5D for parameters roughly corresponding to the human and yeast genomes. We assume that the genome contains *n* = 20000 CREs for humans (or *n* = 6300 for yeast), each *ℓ*= 2000bp (300bp) long. The rest of the genome consists of *L/ℓ*− *n* non-CRE loci, with genome size *L* = 3.2 × 10^9^bp (1.2 × 10^7^bp). We assumed *T* = 1500 (300) TFs, each binding a fraction *f* = 0.1 (0.1) of CREs.

In Fig. 5D, we varied the genome size *L* to show how the number of non-CRE loci affects the information calculations. If TFs bind independently, the information grows with *L*, because binding must be more specific to avoid the additional non-CRE loci. With chromatin, this growth practically disappears: the first, CRE/non-CRE specification term in 10 is negligible. At the actual values of *L*, information savings due to chromatin are very large.

These estimates might be optimistic if some non-CRE loci need to be open in practice, reducing the advantage of chromatin. Also, implementing the additional level of regulation itself requires information: genes involved in chromatin regulation, and their own regulatory elements, must be encoded in the genome. Nonetheless, even if 10^4^ bits were needed to encode each such additional gene, several can be afforded in yeast, and many in humans, from the available savings. This suggests an economy of scale principle in gene regulation: with many genes and TFs, even a small efficiency improvement in encoding all their interactions can pay for encoding a new gene that implements it.

### Biophysical constraints and overspecification

TF binding is thought to be probabilistic and not binary on the timescales relevant for gene regulation. A given target locus can be bound by a TF with a probability that depends on the CRE sequence, TF properties, and its concentration; the resulting gene expression is a function of that binding probability. How does this probabilistic relaxation of the simple binary model of Fig. 2 affect our findings?

Here we extend the simple binary model by postulating that the probability *p*_*B*_ of a TF being bound to the CRE target locus is a function of sequence-dependent binding free energy Δ*G, p*_*B*_ =(1 + *e*^Δ*G*^)^−1^. At face value, this ansatz assumes that the biochemistry of TF-DNA interactions is in thermodynamic equilibrium.

While this is likely the case for prokaryotes [40], it is unlikely to be true in eukaryotes; nevertheless, there, too, the probabilistic (even though non-equilibrium) nature of molecular regulatory interactions is likely to be the norm [41]. Assuming further that a random CRE has Δ*G* drawn from a normal distribution with mean *ζ* and standard deviation *σ*, e.g., because it is the sum of independent free energy contributions from a number of nucleotides (with a continuous approximation for simplicity), we can derive our expectation under neutrality and subsequently ask about selection. To be clear, in this setup, *λ* = {*ζ, σ*} take the role of regulatory parameters that determine the replicated GP map. The corresponding extended model is illustrated in Fig. 6A. The transition from binding to not binding takes place approximately between Δ*G* = − 5 and 5, and depending on *ζ* and *σ*, sequences may exist on either side as well as inside the transition zone.

**Figure 6:**
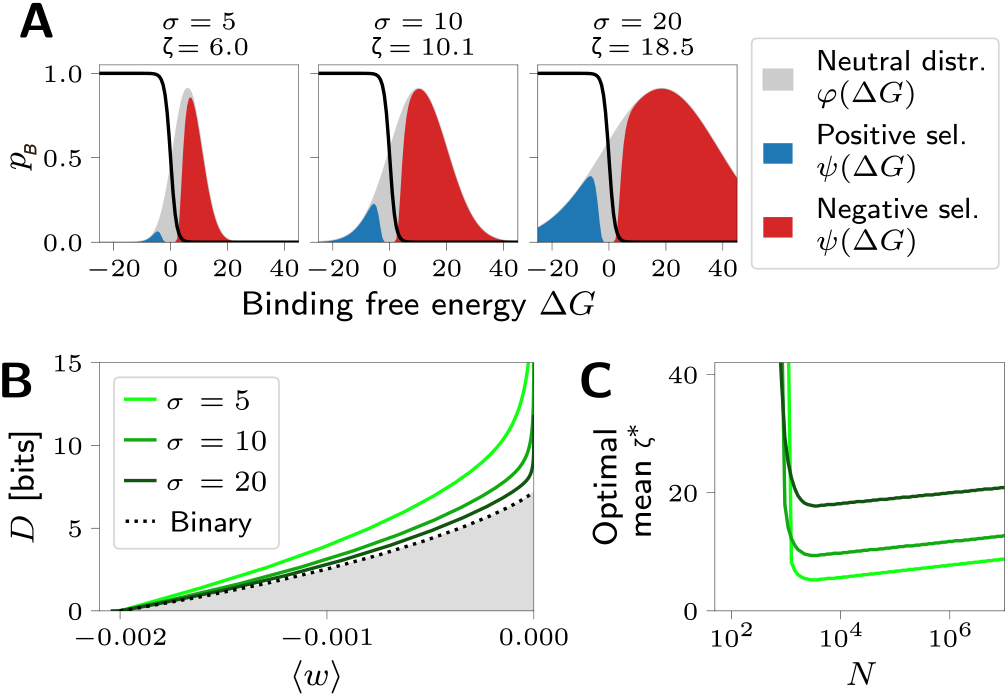
Biophysical constraints lead to overspecification in an extended model with probabilistic TF binding. **(A)** TF occupancy depends on the binding free energy Δ*G*. Under neutrality, Δ*G* in our model is normally distributed with mean *ζ* and standard deviation *σ*; selection shapes individual CREs to drive sufficiently strong (low Δ*G*, blue) or weak (high Δ*G*, red) TF binding. Intermediate Δ*G* (gray transition zone) tend to be avoided. Probability of a TF being bound to the target CRE locus is shown for three choices of regulatory parameters *λ* = {*ζ, σ*} (left-to-right). **(B)** Fitness-information trade-offs achieved by optimal *ζ*^*\**^ with various *σ* (legend). For large *σ*, we approach the binary model, as fewer sequences have intermediate occupancies that need to be avoided. “Overspecification” refers to the extra information needed at fixed ⟨*w*⟩ because of biophysical constraints (i.e., the difference between the green and dotted black curves). **(C)** Values of *ζ*^*\**^ that optimize free fitness as function of *N*, for three different values of *σ* (green color shade, legend in B).

Selection acts on the binding phenotype, as before: but now, the log-fitness penalty is *p*_*B*_*s* for spurious binding (FP) and (1 − *p*_*B*_)*s* for failing to bind when needed (FN). Either type of selection on individual CRE will restrict the Δ*G* distribution to sufficiently high or sufficiently low values (red and blue distributions in Fig. 6A). Importantly, sequences inside the gray transition zone will be eliminated by positive and negative selection alike. Intermediate binding, such as at *p*_*B*_ = 0.5, is never desirable in this model, and an IT-efficient frequency-matching code would not assign it any genotypes. Yet with a unimodal distribution over Δ*G*, it is impossible to avoid intermediate binding without also eliminating either of the two desirable phenotypes (*p*_*B*_ ≈ 1 and *p*_*B*_ ≈ 0). The next best solution would be to make *σ* very large, so that the vast majority of genotypes is far on either side of the transition. But *σ* is very likely biophysically constrained, for instance, by the strength of molecular recognition interactions (set by the energetics of the hydrogen bonds) or the binding site length (set by the physical footprint of the TF protein), so that this limiting solution may likewise be inaccessible in practice.

Biophysical constraints mean that achieving the expected log-fitness takes more information compared to when the constraints are not acting, as shown in Fig. 6B. The optimal *ζ*^*\**^ behaves similarly to the parameter *q* of the simple binary model (Fig. 6C). In small populations, CREs cannot adapt anyway and the best choice is to accommodate the majority of CREs (which require no binding), by choosing a very large *ζ*. In large populations, intermediate values of *ζ* can approximate frequency matching. But this approximation falls short of the IT-efficient solution due to the sequences in the transition zone, necessitating extra work to be done by selection acting individually, at every CRE target locus, so as to deliver a functional phenotype. Taken together, this is precisely an instance of “overspecification” [42] – due to constraints, selection must accumulate and maintain more information *D* to support function than a straightforward and model-independent combinatorial analysis would suggest.

### Robustness and the strong mutation regime

A major assumption of the theory presented thus far is that evolution proceeds by successive fixations in otherwise monomorphic populations (“the fixed states approximation”). While this assumption may be justified for individual target loci with *L*∼ 200 bp for human populations and marginally justified for, e.g., *Drosophila*, it is certainly not satisfied when considering all coevolving CREs and their regulator loci on a replicated GP map. Because there may be genetic variation within individual target CRE loci, and because our setup crucially depends on the epistasis between regulatory loci and a large number of target loci (Fig. 1), we now turn our attention to the strong mutation regime.

This regime is technically more difficult to study. The level of description must move from distributions over individual genotypes, *ψ*_*i*_, to distributions over “population states”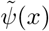, defined by genotype frequencies *x*, in diverse populations [23]. Fortunately the free fitness theory was originally developed and later extended with this case in mind [20, 21, 43]. In analogy to the monomorphic case, one can show that the distribution over population states maximizes population-level free fitness (SI Appendix), a balance between log mean fitness and an information term 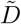, which upper bounds genetic information 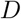, i.e., 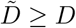.

The difference between the two information measures 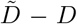 vanishes when mutation is weak and the population is mostly monomorphic (SI Appendix). In that limit, one recovers the free fitness formula of 3, and, consequently, our entire theoretical framework under the fixed states approximation. Taken together, even in diverse populations selection optimizes regulatory parameters by maximizing free fitness – but there are two key differences with fixed states approximation in the weak mutation limit.

First, at the population level, free fitness results do not automatically translate into a distribution over regulatory parameter values *λ*, as they did in 5. This is because *λ* itself can have diverse values in a population, depending on the genotype frequencies at regulatory loci. Optimization still takes place, but in the space of population states, each with some variation in *λ*. If selection is strong and *λ* impacts many target loci, states with too much diversity in *λ* may be suppressed, but this can cause departures from 5, by favoring values of *λ* that are more robust to mutational changes, or values where such changes cause smaller reductions in free fitness.

Second, mutational robustness is also important at the target loci. In the weak mutation regime under the fixed states approximation, free fitness favored maximizing the number of fit genotypes. In the strong mutation regime, the focus shifts from genotypes to populations. Populations are selected towards high log-mean-fitness, while minimizing the KL divergence from the neutral distribution over population states, 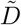. But now 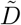 contains an additional term (SI Appendix), which reflects the effects of selection on genetic variation. This term depends on the mutational robustness of the occupied genotypes. If the genotype *i* is robust, mutations to it will have little effect on fitness; consequently, the population is likely to have the same levels of diversity under selection as under neutrality, and the extra term contribution to 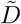 will be low. On the other hand, if *i* is not robust, e.g., because it sits at a sharp fitness peak or near a steep cliff, the inferior mutational neighbors are likely to be missing from the population under selection, even though they would have been present under neutrality. Minimization of 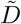 therefore automatically favors robust genotypes, which, *per se*, is not the case for minimization of *D*.

We illustrate the effects of strong mutation in Fig. 7. To explore the interplay between genetic information, fitness, and robustness, we extend our simple binary model of Fig. 2 yet again, this time by introducing a stylized mechanism for TF-DNA molecular recognition. In this extension, a TF that recognizes *ℓ*= 20 bp sites binds to the target CRE locus, here for simplicity treated also as an *ℓ*bp sequence, if there are ≤ 8 mismatches between the TF motif and the CRE; this setup keeps the binding/no-binding nature of the simple binary model, but formulates it in sequence space. Some target loci are under selection to bind the TF and the remainder under selection against binding, as before. Individual-based asexual Wright-Fisher simulations with point mutations highlight two main effects of the increased mutation rate. First, as the mutation rate *u* grows, Fig. 7A shows that selection is increasingly incapable of maintaining CRE target loci that should bind the TF cleanly separated from loci that should not; this expected result, in line with the classic literature on mutation load, suggests that high fitness solutions get increasingly difficult to realize at higher mutation rates. This is also reflected in Fig. 7C, where the best achievable log-mean-fitness (even as *N*→ ∞) at high *u* falls below the values achievable at lower *u*.

**Figure 7:**
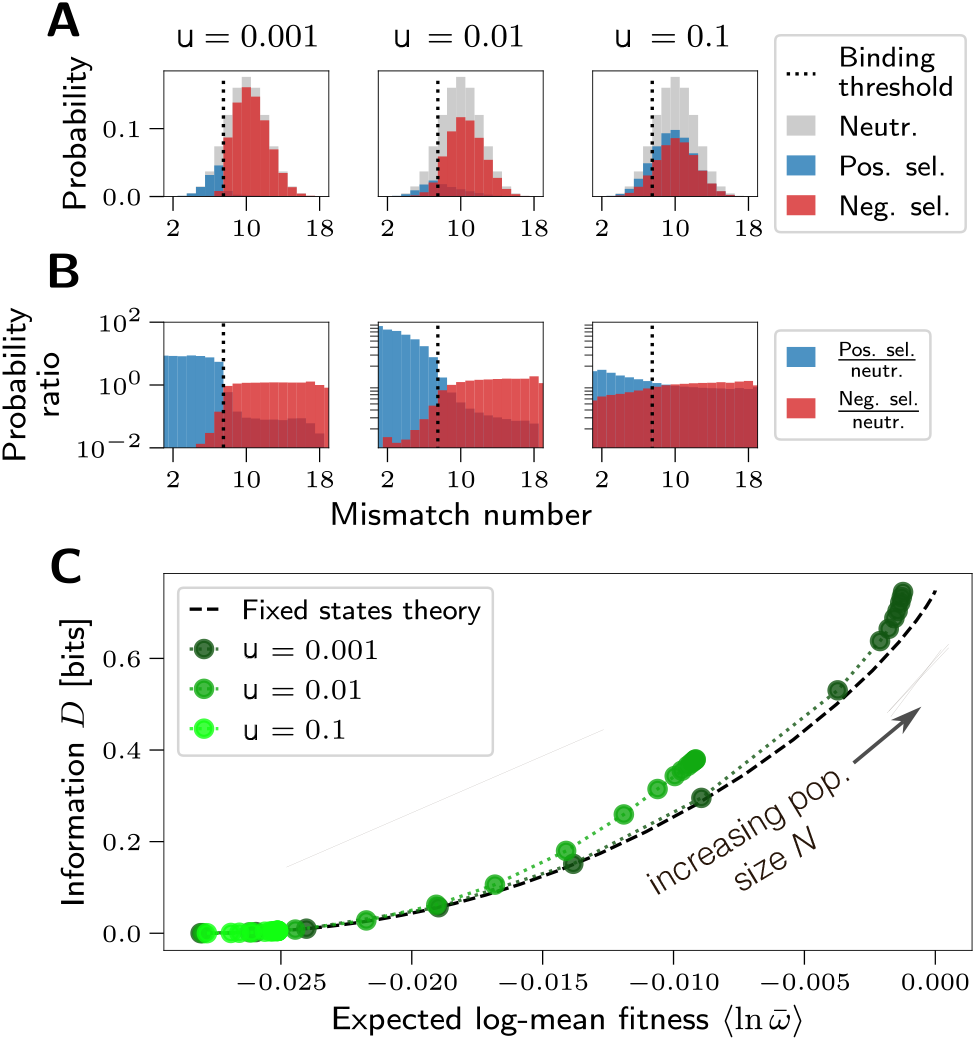
Strong mutation hinders adaptation but promotes mutational robustness. Individual-based Wright-Fisher simulations of *ℓ*bp CRE evolution, where each CRE can only harbor a single TF binding site (see text). TF binds if the binding site contains less mismatches with TF’s preferred motif than the “binding threshold”. *u* is the per-bp per-generation per-individual mutation rate. **(A)** Mismatch distributions under neutrality (gray) and under selection for/against binding (blue/red), as mutation rate *u* increases (left-to-right). **(B)** Ratios of mismatch distributions under selection vs. neutrality. **(C)** Mean-log-fitness vs. information trade-off plot, parameterized by the population size, *N* (arrow direction). At high mutation (bright green), high fitness values cannot be achieved even at large *N*, due to mutation load. When a given log-mean-fitness can be achieved, strong mutation scenarios (green shades) require more information *D* compared to the fixed states approximation (black dashed): this is robustness overspecification. Fixed parameters: *n* = 10 CREs of *ℓ*= 20 bp each, fraction *f* = 0.2 under positive selection and 1 − *f* under negative selection, *s* = 0.1 log-fitness penalty per error at each of the *n* target loci, population size *N* = 25 (except in C, where *N* is varied).

Fig. 7B zeros in on a more subtle effect. Under the fixed states approximation, the equilibrium theory predicts that the ratio of mismatch distributions at target loci selected for or against binding should solely be a function of the corresponding fitness – i.e., this ratio should be constant (flat) above and below the mismatch threshold, with a sharp step in between. Yet as mutation rate increases, we observe strong deviations from this expected constancy (Fig. 7B), to favor genotypes that are further away from the threshold, in a clear signature of an automatic selection for mutational robustness. Ultimately, strong mutation modifies the theoretical trade-off between the log mean fitness and information *D* (since *D* is no longer equal to 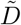 at high *u*), as shown in Fig. 7C: assuming that the particular expected log-mean-fitness is achievable at a given mutation rate *u*, selection must accumulate more information *D* at higher *u*, as it concentrates population states around genotypes that are both fit as well as mutationally robust. This can be interpreted as a case of robustness-associated overspecification. Mutational robustness is a recognized factor in evolution in general [44–46], and in the context of gene regulation in particular [47, 48]. Importantly, it is intrinsically a local phenomenon on a GP map, associated with the preference for genotypes with fit mutational neighbors [46]. While low information *D* also prefers GP maps with a large number of fit genotypes and can be seen as a kind of “genotypic redundancy” [49], *D* does not care how these genotypes are arranged in sequence space, and in particular, whether they are mutational neighbors to each other or not. The relationship between mutational robustness and *D* is therefore complicated. On the one hand, mutational robustness may emerge indirectly, as a by-product of the minimization of *D* and irrespective of the mutation rate, if the genotypes with high fitness happen to be close in sequence space, perhaps because of biophysical constraints [48, 50]. On the other hand, when mutation rate is high, robustness becomes important in its own right, and will be preferred as population-level free fitness increases and 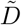 is minimized. Further theory work is needed to disambiguate the two effects, to quantify robustness within the information-theoretic framework, and to simulate replicated GP maps where the arrangement of fit solutions in sequence space can also evolve. In general, however, we hypothesize that at strong mutation, the regulatory parameters *λ* would become mutationally robust themselves, as well as associate high fitness with many mutationally robust genotypes at the target loci.

## Discussion

In this paper we used the free-fitness theory and the recent developments of the genetic information formalism [23, 29] to explore the genotype-phenotype maps for gene regulation.

The genetic information that enters our calculations is similar to the bioinformatics concept of the information content (IC) of TF binding motifs [28, 51]. IC quantifies, for a collection of binding sites, how different they are statistically from the rest of the genome (a comparison similar to that between distributions under selection vs. under neutrality, if selection alone drove the evolution of the binding sites) [52]. The requirement to localize TF binding to a unique site within the genome implies a necessary amount of IC. Here, we asked a similar question more generally, and about any regulatory task. How much information does a CRE need to contain so that it interacts with a given set of TFs, or expresses the gene in a given set of environments?

We showed that equilibrium population genetics leads to an optimization principle for regulatory parameters: they will be driven to values that maximize the fraction of random sequences encoding high-fitness phenotypes. An analogy with coding in information theory suggests that, under strong selection, “frequency matching” will ensure that most of the sequence space is devoted to phenotypes required at many loci, and a lot of genetic information will be needed at loci that require unusual phenotypes. In ideal circumstances, the information needed in an IT-efficient GP map is given by the entropy associated with phenotype requirements across the genome. This can be increased due to biophysical constraints or the need for robustness in high-mutation regimes.

We highlight five key implications of our theory.

**First**, optimization of regulatory parameters and, consequently, of replicated GP maps, can take place also if selection at each target locus is weak compared to drift, so long as the number of target loci is large. This novel route to evolutionary optimization can be used to make *ab initio* predictions about certain properties of GP maps even without knowing the molecular mechanisms that underpin their structure. In this context, being optimal implies a GP map on which selection has to exert least “work” by accumulating a minimum of information *D* in target loci to ensure function and maximize fitness. Optimal regulatory parameters may depend on selection strength (*Ns*), but will be driven to their maxima regardless of the *Ns* value itself.

In our toy model, fitness multiplies over loci, and so if selection is weak relative to drift at each locus, each CRE is poorly adapted, as is the whole system. Lynch et al. [25, 53] emphasize this as a “drift barrier” to molecular adaptation. However, the whole system may still function reliably, even if random drift dominates the evolution of its individual components. In the companion paper, we consider multiple weak binding sites within a CRE, which together give reliable regulation [17]. More generally, such situations can be represented by suitable nonlinear maps between net binding and fitness. This is just the regime of the infinitesimal model in quantitative genetics, in which phenotypic traits adapt predictably, even while selection hardly perturbs the underlying loci from neutrality [54].

**Second**, our theory makes clear that considering common, even if (at face value) biologically uninteresting phenotypes, is essential to understand the optimal – and likely evolved – structure of GP maps. While we typically focus on functional and rare outcomes, e.g., by asking “In which genomic regions does a particular TF bind?”, our arguments show that the major determinant of an optimal GP map lies with the question “In which genomic regions should this TF not bind?” Recent work clarifies this point in the systems biology context, by studying the constraints due to “deleterious crosstalk”, or misregulation of gene expression caused by off-target, non-cognate binding of TFs to CREs (which mimics our per-locus selective penalty *s* for erroneous binding) [14, 55, 56]. Crosstalk and binding promiscuity have also been implicated for the evolvability of regulatory sequences [1, 50, 57]. Here, and especially in the companion paper [17], we demonstrate the essential role of crosstalk and selection against binding in shaping optimal genotype-phenotype maps.

**Third**, we find that the difference in free fitness between optimal and sub-optimal GP maps can be dramatic, amounting to millions of bits across the whole genome for plausible parameters, due to the sheer number of TF–DNA interactions affected. The magnitude of such savings is critical, since they can pay for novel regulatory mechanisms which themselves must be genetically encoded. Using a toy model, we suggest this to be a possible reason for the evolution and maintenance of chromatin-based gene silencing. More generally, we demonstrate that genetic information provides a quantitative framework to evaluate, compare, reason about, and perhaps even predict the existence and evolutionary role of putative regulatory mechanisms.

**Fourth**, optimal GP maps, in the strict information-theoretic sense defined here, share many properties with maps that should be very evolvable [58]. If we consider *de novo* evolutionary adaptation, which starts at a random point in sequence space, then maximizing the fraction of fit sequences (i.e., minimizing genetic information *D*) should correlate with minimizing the time needed for the adaptive walk to locate and fix a fit solution [29]. While the theory presented here is clearly an equilibrium theory, it suggests a dynamical extension, termed “optimize-to-adapt,” that we pursue in detail in the companion paper [17]. In this simulation-based approach, regulatory parameters are optimized numerically (as predicted by our equilibrium theory, but assuming that it happens over a very slow timescale) so as to speed up the adaptation of CRE sequences, which can be simulated explicitly and is assumed to happen faster than the regulatory parameter evolution. As a result, we can define evolvable maps as those whose regulatory parameters lead to highest *de novo* adaptation rates for functional CREs. We show that there is an excellent correspondence between the steady state, information/free-fitness-based theory derived here, and the dynamical instantiation studied in the companion paper.

**Lastly**, the presented theory allows us to reason more broadly about information accumulated in genomes. Quantification of the necessary information is very pertinent in the context of debates about genome functionality. Comparative genomics suggests that up to about 15% of the human DNA is under detectable selection [59,60], most of which is non-coding and must affect fitness via regulatory processes. But a much larger fraction (up to about 80%) of the genome seems to have the capacity for, or shows signs of involvement in, such regulatory processes [5, 61–63]. This has led to debates about technical problems and underappreciated evidence [64], the meaning of function [65–67], and insights from population genetics theory [68, 69]. One source of difficulty is the unknown and possibly complicated genotype-phenotype relationship for regulatory sequences. Long regions of DNA could perform simple functions that only weakly constrain their evolution. Quantification of the information necessary for regulatory tasks, as we propose here, avoids the dichotomy between functional and non-functional; furthermore, it can be performed without a complete knowledge of empirical GP maps or molecular mechanisms involved, by focusing on tractable computations over GP maps that are theoretically optimal for a given regulatory task.

We briefly comment on some of the limitations of our theory. First, the focus on stationary distributions may be problematic since approaching them might be too slow, e.g., because selection changes on a faster time scale or because populations might settle into local optima and never explore more of the sequence space. While this is indeed a serious limitation, our results might be nonetheless relevant. TF binding sites are short and the difference between a functional binding site and random sequence is often only a few point mutations [7, 8], making exploration of the sequence space possible at least for individual regulatory elements. Also, even if the equilibrium is not reached, the information needed to encode a phenotype could be either a useful statistic or a theoretical measure that predicts well the rates for evolutionary emergence of novel functions [17].

Second, the existence of equilibrium distributions requires detailed balance. At low mutation, this restricts the form of mutation rates, excluding structural mutations such as duplications or deletions. At high mutation, it further restricts us to scenarios with linkage equilibrium between small loci (with negligible standing variation at each), excluding also finite recombination rate. What happens outside these regimes is an open question. Structural mutation that produces repetitive sequences could, for example, lead to optimal GP maps that employ such sequences for regulatory tasks, as seems to be the case with transposable elements [70]. Non-equilibrium statistical physics methods might be used to study these technically difficult cases.

Lastly, an important empirical difficulty that hampers the broader application of our framework is the assumed knowledge of what function is being selected for. If we knew what regulatory phenotypes are required along the genome, we could predict the optimal GP maps and compute the necessary information – but the requirements are actually unknown. Instead, we typically make assumptions about selection based on what is observed. While the first steps in bringing together optimization theories with inference have been made [34], further work is required to bring our framework into direct contact with genome-scale data.

## Acknowledgments

Michal Hledik (Institute of Science and Technology Austria) co-designed this research, performed the research, and wrote the first manuscript draft. This research also forms a part of his thesis work, *Genetic information and biological optimization*, deposited at DOI: 10.15479/at:ista:15020. Based on his contribution, M. Hledik should be the first author on this paper, but because we could not get in touch to obtain consent to deposit, biorxiv policy dictates that we acknowledge his contribution here. GT and MH acknowledge the support of Human Frontiers Science Program grant RGP0034/2018. NB acknowledges support from ERC grant Haplo-typeStructure 101055327.

## Notes

### Competing Interest Statement

The authors have declared no competing interest.

