## Supplemental Info for "Evolution and information content of optimal gene regulatory architectures"

Michal Hledík, Nicholas H Barton, Gašper Tkačik

June 10, 2025

In this short SI Appendix, we outline how the information theoretic approach can be applied also beyond the "fixed states approximation," to regimes where the mutation rate is high enough such that populations are genetically diverse. The derivations closely mimic those outlined in the main paper which is why we do not reproduce them there. There are, however, certain important definitions and terms that are new and unique in the high mutation rate regime, including the emergence of terms that relate to mutational robustness.

We mostly follow the notation of Ref. [2], with the primary goal to point out the interesting phenomenology, rather than to be systematic and verbose.

On the population level, the equilibrium distribution for diverse populations, $\tilde{\psi}(x)$, is defined over a vector of genotype frequencies $x$. In principle, these could be complete genotype frequencies with no recombination, or allele frequencies at uncorrelated sites. The equilibrium distribution $\tilde{\psi}(x)$ then also takes a Boltzmann form

$$\tilde{\psi}(x) \propto \tilde{\varphi}(x) e^{2N \ln \bar{\omega}(x)}. \quad (1)$$

$\tilde{\varphi}(x)$ is the neutral distribution, and $\bar{\omega}(x) = \sum_i x_i \omega_i$ is the mean fitness within a population, in which genotypes $i$ come with frequencies $x_i$ and associated genotypic fitness values $\omega_i$. The distribution $\tilde{\psi}$ maximizes population-level free fitness,

$$\tilde{F} = \langle \ln \bar{\omega} \rangle - \frac{1}{2N} \tilde{D}. \quad (2)$$

In analogy to $D$ of Eq. (2) in the main paper, $\tilde{D}$ is likewise a type of genetic information introduced in Ref. [2], but applicable to diverse populations: it quantifies how different the distribution over population states is under selection, $\tilde{\psi}(x)$, vs under neutrality, $\tilde{\varphi}(x)$. We can connect the two information quantities by computing the probability that a genotype $i$ is observed by randomly sampling it from a population whose genotype frequencies are given by $x$:

$$\psi_i = \int \tilde{\psi}(x) x_i dx \qquad \text{and} \qquad \varphi_i = \int \tilde{\varphi}(x) x_i dx. \quad (3)$$

It can be shown rigorously that the population-level information $\tilde{D}$ is always larger or equal to the genotype-level information $D$ [2],

$$\tilde{D} = D + \sum_i \psi_i \int \tilde{\psi}(x|i) \ln \frac{\tilde{\psi}(x|i)}{\tilde{\varphi}(x|i)} = D + D(X|G) \geq D \quad (4)$$

where $\tilde{\psi}(x|i) = \tilde{\psi}(x)x_i/\psi_i$ is the conditional distribution over population states given that a randomly sampled genotype was $i$. The inequality holds because the second term, $D(X|G)$, is a non-negative conditional Kullback-Leibler (KL) divergence [1]. This non-negative term, $D(X|G) = \tilde{D} - D$, that we refer to in the main text as the "extra term", captures the effects of selection on the *variation* in the population.

The divergence $D(X|G)$ vanishes and $\tilde{D} = D$ when mutation is weak and the population is mostly monomorphic. In that limit, once a single genotype $i$ is sampled, one can be practically certain that it is fixed in the population regardless of selection (i.e., $\tilde{\psi}(x|i) = \tilde{\varphi}(x|i)$ and both peak at $x_i = 1$). Furthermore, the log-mean-fitness, $\ln \bar{\omega}(x)$, can be replaced with the log-fitness $w_i$ of the fixed genotype $i$, thereby recovering the free fitness formula of (**??**), and, consequently, our entire theoretical framework under the fixed states approximation.

As in Eq. (4) of the main paper, we consider a joint system of regulatory loci and their targets, to compute the marginal distribution over the genotype frequencies $x^R$ at the regulatory loci,

$$\tilde{\psi}(x^R) \propto \tilde{\varphi}(x^R)e^{2N\tilde{F}(x^R)}, \tag{5}$$

where $\tilde{F}(x^R)$ is the population-level free fitness of the target loci conditional on the regulator loci. Therefore, even in diverse populations, selection acts to optimize regulatory parameters by maximizing free fitness – in the main text we discuss, however, the two key differences between the monomorphic and diverse settings.

In summary, when mutation is strong, the trade-off between (log-) fitness and genotype-level information $D$ changes to a tradeoff between (log-mean-) fitness and population-level information $\tilde{D}$. Eq. (4) suggests that the latter could be decomposed into $D$ plus a term associated with robustness-associated overspecification. While clarifying these relationships will require further theory work, we already get a glimpse of understanding evolution in genetically diverse populations in terms of a triple tradeoff between fitness, information, and mutational robustness. Our current view suggests that regulatory parameters $\lambda$ of a replicated GP map will evolve to become mutationally robust themselves, as well as to associate high fitness with many mutationally robust genotypes at the target loci.
